# How beat perception coopts motor neurophysiology

**DOI:** 10.1101/805838

**Authors:** Jonathan J. Cannon, Aniruddh D. Patel

**Author notes:** Submitted to *Trends in Cognitive Sciences* (revision 2).

## Abstract

Beat perception is central to music cognition. The motor system is involved in beat perception, even in the absence of movement, yet current frameworks for modeling beat perception do not strongly engage with the motor system’s neurocomputational properties. We believe fundamental progress on modeling beat perception requires a synthesis between cognitive science and motor neuroscience, yielding predictions to guide research. Success on this front would be a landmark in the study of how “embodied cognition” is implemented in brain activity. We illustrate this approach by proposing specific roles for two key motor brain structures (the supplementary motor area, and the dorsal striatum of the basal ganglia) in covert beat maintenance, building on current research on their role in actual movement.

**Highlights:**

⍰ Components of the brain’s motor system are activated by the perception of a musical beat, even in the absence of movement, and may play an important role in beat-based temporal prediction.
⍰ Two key brain regions involved in movement, the supplementary motor area and dorsal striatum, have neurocomputational properties that lend themselves to beat perception.
⍰ In supplementary motor area, neural firing rates represent the phase of cyclic sensorimotor processes.
⍰ Supplementary motor area’s involvement in perceptual suppression of self-generated sounds suggests that it could play a broader role in informing auditory expectations.
⍰ Dorsal striatum plays a central role in initiating and sequencing units of movement, and may serve similar functions in structuring beat-based temporal anticipation.

## What is the motor system doing during beat perception?

Human auditory perception extends beyond what is presented to our senses. When an auditory rhythm has induced the sense of an **isochronous**(see Glossary) beat, a listener implicitly predicts where each beat falls in time, even though some beats may not be marked by sound and some sounds can occur at non-beat times (see Figure 1A and supplemental sound file). In humans, this **beat maintenance**is closely related to movement: a perceived beat often elicits rhythmic motion [1–3], and rhythmic motion improves accuracy on beat-based perception tasks [4,5]. Further, **fMRI**(functional magnetic resonance imaging) reveals that regions of the motor system are active and interacting while a beat is perceived even in the absence of movement [6–8]. The most consistent of these are the **supplementary motor area (SMA)**of cortex and the dorsal section of the **striatum**of the subcortical basal ganglia including the **putamen**[6–12], which are parts of the brain’s “direct pathway” for motor control (Figure 1B).

**Figure 1:**
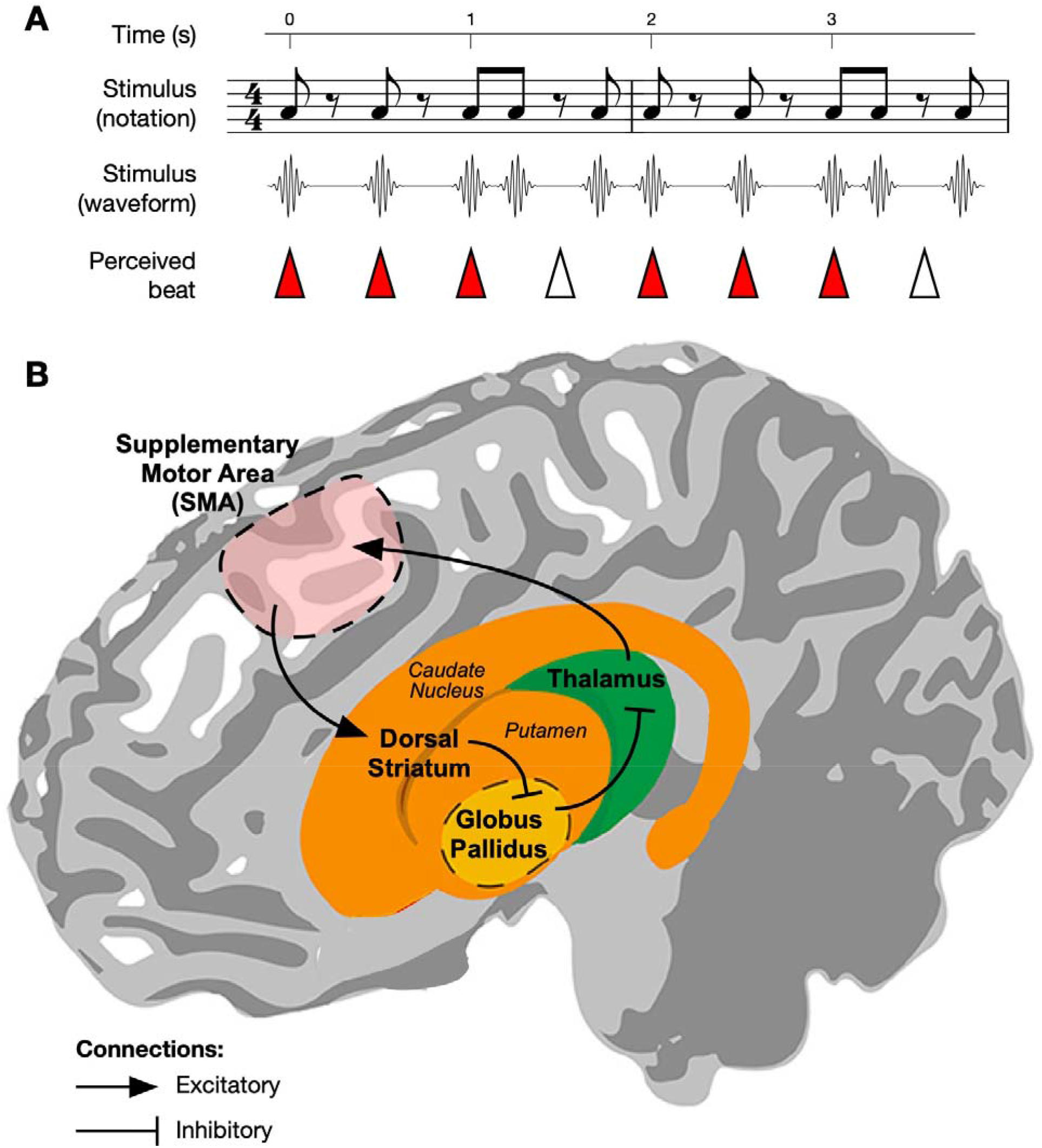
Beat perception and the motor system. **A)** A simple, repeating non-isochronous rhythm is shown in music notation and illustrated as an acoustic waveform. The rhythm induces an isochronous beat percept (beat locations shown by triangle tips). Note that some beats are not marked by sounds (hollow triangles), and some sounds occur at non-beat times. The reader is encouraged to listen to the corresponding sound example (in which the above rhythm is looped several times) to verify the beat percept. **B)** A simplified schematic of the direct pathway loop connecting the two key areas discussed in this paper in reference to beat perception. The supplementary motor area (SMA) projects to the dorsal striatum (consisting of the putamen and caudate nucleus). Dorsal striatum inhibits the internal segment of globus pallidus (shown here hidden behind the striatum), which inhibits thalamus. Closing the loop, thalamus excites SMA. During physical movement, this pathway is thought to selectively disinhibit populations in SMA necessary for the next sub-movement in a sequence. Here, we propose that the same process is used to disinhibit populations that generate the temporal dynamics necessary to covertly anticipate the next beat in a rhythm. Figure adapted from [116]. CC BY 3.0 https://creativecommons.org/licenses/bv/3.0/

Although the motor system’s activation by covert beat maintenance could be a passive byproduct of the frequent association of beat perception and movement, a number of authors have argued that the motor system plays a causal role in covert beat perception [11,13–16]. Some have specifically suggested that beat-based temporal predictions arise from “simulated rhythmic actions” in the motor system [17–21], For example, the “Action Simulation for Auditory Prediction”**(ASAP)**hypothesis [19] posits that simulated actions in motor planning regions provide temporal predictions about beat times that influence activity in auditory regions. However, this hypothesis currently lacks a neurophysiological account of the “simulated action” and its relationship to actual movement.

We argue that the idea of temporal computations via internal actions can be mapped on to a specific proposal about the roles of the SMA and dorsal striatum in beat maintenance. We focus on the roles of these structures because we find that an understanding of their motor functions can be generalized to account for the maintenance of an internal pulse underlying a rhythm. Crucially, this proposal draws heavily on known aspects of motor system physiology, thus integrating cognitive science and motor neuroscience.

## The ASAP hypothesis and action simulation

Beat perception involves spontaneously recognizing that a rhythm supports a beat and inferring its tempo and phase **(beat inference)**, and a subsequent process of beat maintenance (see [22,23] for neural research dissociating these processes). The ASAP hypothesis establishes a general framework for understanding beat maintenance in the brain. It posits that auditory cortex sends auditory timing information to motor planning areas, which initiate simulated actions timed to terminate on the next beat. These motor signals travel to auditory cortex via temporally-precise reentrant signaling [24], where they predict beat timing and shape the perceptual interpretation of rhythms. ASAP argues that this co-opting of motor planning activity for auditory prediction is effective because the motor system is an excellent generator of temporal patterns on the time scale of musical beats. Our strongest sense of beat occurs when inter-beat-intervals are between ~4OO-12OO ms, and humans show a preference for beat periods around 600 ms [25], near a natural frequency of human locomotion [26].

ASAP proposes that communication between motor and auditory areas in beat perception relies on a dorsal auditory pathway that bidirectionally links the premotor, parietal, and temporal cortices [27] (see [28] for a test of this proposal). It further proposes that the temporo-parietal part of this pathway was strengthened in humans (relative to other primates) over evolutionary time as the human lineage became vocal learners [29,30]. The strength of this pathway in humans could help explain why auditory rhythms so readily engage the motor system in our species. The pathway’s relative weakness in monkeys [31,32] could account for why they (unlike humans) do not spontaneously move predictively when synchronizing movements to periodic rhythms, yet do single-interval timing in a way that is behaviorally and neurally similar to humans [33,34],

However, an essential aspect of ASAP remains conceptually underdeveloped: the simulated action. Up to now, ASAP has remained agnostic as to how a simulated action is neurally instantiated in the motor system. Here we propose that the simulated actions proposed by ASAP consist of precisely patterned neural time-keeping activity in SMA, orchestrated and sequenced by activity in the dorsal striatum. Activity in SMA is tuned to play out over the course of one beat period, informing auditory anticipation over the course of each inter-beat interval and adjusting its state to continuing auditory cues. Dorsal striatum uses contextual input from motor and sensory cortices to select and disinhibit consecutive units of SMA activity (which we call **proto-actions).**Below, we build up these hypotheses in detail and review the evidence supporting them.

## SMA in beat maintenance

In humans, the supplementary motor area (SMA) is activated by a range of beat-based tasks, including self-paced finger tapping, tapping entrained to music, and (importantly for this discussion) tasks involving the covert tracking of a beat [6-8,35]. Differences in SMA activation account for individual differences in performance on covert beat-based timing tasks [36,37], suggesting that SMA plays a functional role in covert beat generation. A covert repetitive timing task in monkeys has demonstrated that neural activity in SMA can maintain an ongoing “metronome,” tracking task-relevant periodic time intervals in the absence of sensory cues [38].

### “Proto-action” for timing and prediction

As we formulate hypotheses of the roles played by SMA in beat-based predictive timing, the ASAP hypothesis suggests that we look to its role in movement. To appreciate this role, it is important to consider motor control through the increasingly influential **dynamical systems**lens [39,40]. While the classical “tuning/decoding” perspective asks how activity of specific neurons may represent different aspects of movement (such as direction, velocity, etc.), the dynamical systems perspective asks how interactions between neurons can give rise to reliable, flexible time series of motor commands. For example, monkey electrophysiology and computational modeling have shown that arm reaching is orchestrated by temporal fluctuations in firing rates of neurons in motor cortex, and that during a reach these fluctuations are likely attributable largely to interactions between motor cortical neurons via recurrent connections [40]. These fluctuations can be viewed as tracing out reliable paths (“trajectories”) in an underlying space of possible firing rates, with each position in this space corresponding to a set of signals being relayed to the muscles (see Figure 2 for illustration).

**Figure 2:**
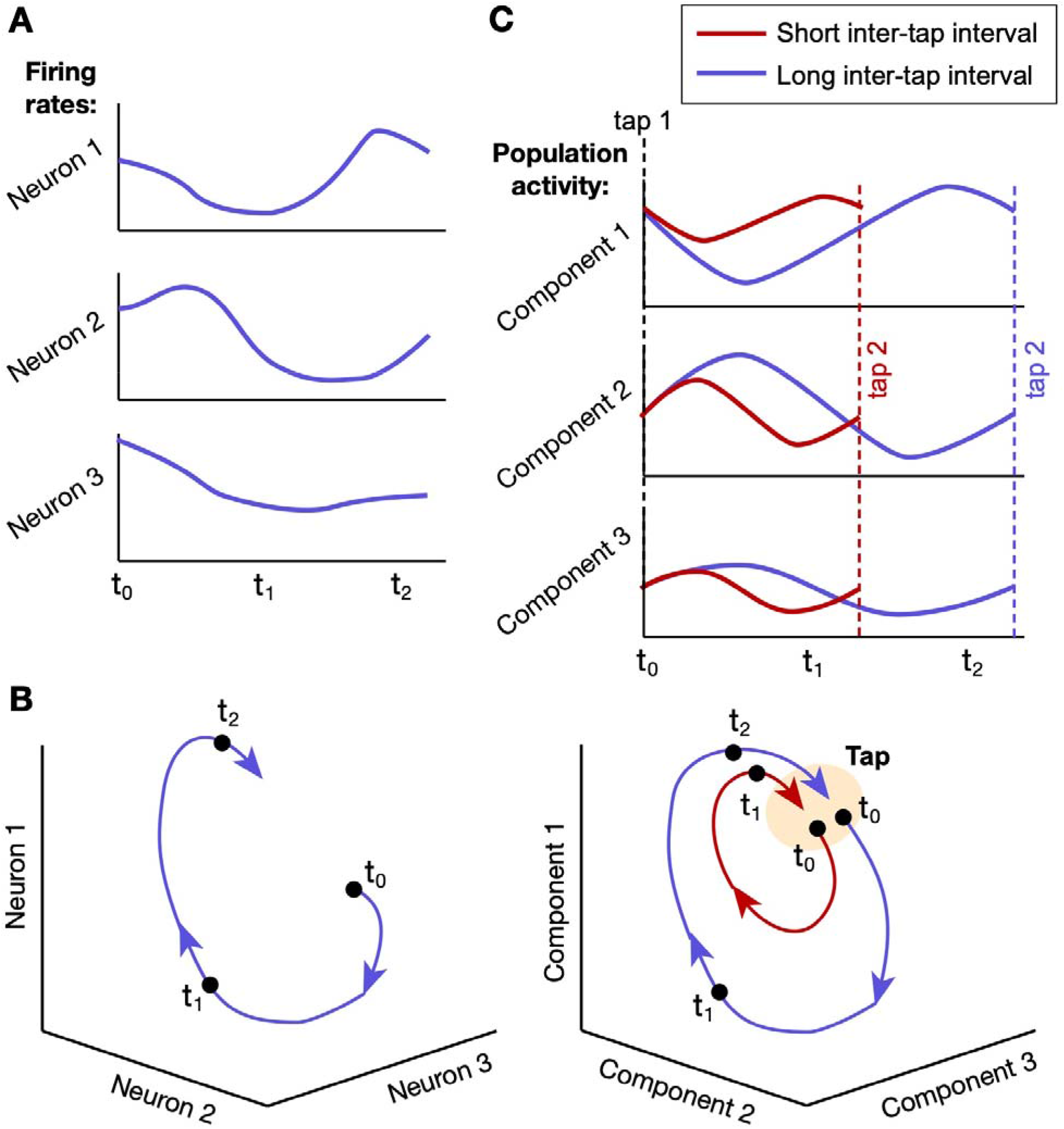
Illustration of time-keeping by a neural firing rate trajectory. In monkey SMA, reliably patterned fluctuations in the collective firing rates of neural populations keep time during deliberately timed behaviors like rhythmic tapping. These fluctuations can be visualized as tracing out paths (“trajectories”) in a high-dimensional space of firing rates, with the shape and time course of the path dependent on the time course of the behavior. A) Fluctuations in the firing rates of three neurons graphed over time during a timed behavior. B) The same fluctuations visualized as a trajectory in a three-dimensional space, where each axis represents the firing rate of one neuron and each point on the trajectory represents the three firing rates at one point in time. Three example time points are marked. C) Trajectories traced by firing rates in large neural populations (i.e., hundreds, rather than just three) are easier to visually understand in “principal component” space [117], where each axis is a “component” corresponding to the correlated activation of many neurons, and where components accounting for the most combined variance in population firing rates are displayed. Empirical research on periodic tapping in monkeys shows that the firing-rate fluctuations in SMA between consecutive taps trace out circular trajectories in principal component space. For shorter inter-tap intervals, firing rates fluctuate faster but with lower amplitudes, creating smaller loops that repeat with a shorter period [46]. (Image adapted from the results of [46] and simplified for illustration.)

Research shows that during movement execution in non-human primates, dynamics in SMA resemble those in motor cortex: neural firing rates during a specific movement pattern follow reliable, repeatable trajectories. But recent results indicate that firing rates in SMA give a more abstract representation of progress through a given action, continuing to change over time through periods of motionlessness (e.g., between consecutive finger taps [41]) and occupying distinct states on consecutive repetitions of the same movement [42–44], Importantly, when motionless behavioral intervals are deliberately lengthened or shortened to meet task demands, some combination of the speed and amplitude of these trajectories and the number of neurons involved adjust accordingly, tuning the system to arrive at a similar end state associated with a behavioral response (e.g., a tap) in a different amount of time [41,45,46] (see Figure 2C). This behavior differentiates the population activity from a strictly “absolute” timekeeper, which would progress similarly regardless of the target interval (and which is well-suited to single-interval timing). Instead, its state offers “relative” timing information about the elapsed time relative to the target interval.

Following ASAP’s analogy of covert inter-beat interval timing to action, we hypothesize that in humans, SMA behaves in covert beat-based timing like it behaves during overt cyclic movement in monkeys (such as repetitive reaching or tapping), but without the accompanying motor cortex activation directing physical action. In other words, we propose that SMA tracks progress through each beat interval by generating firing rate dynamics that are tuned to arrive at a characteristic state at the next beat time, whether or not there is a behavioral response attached to that beat. As noted above, strong auditory-motor connections in humans permit this motor timing information to reach auditory cortex in a temporally precise fashion, enabling beat-based auditory prediction in humans to a degree unmatched by non-human primates.

How is SMA tuned to create dynamics that anticipate the appropriate interval? Analysis of brain activity in regions upstream and downstream from SMA in monkeys (and supporting models) indicate that interval-specific adjustments to SMA trajectories seem to be made through modulation of **tonic inputs**to and/or initial state of an SMA network, with the resulting dynamics generated through local interactions within SMA [45,47,48]. (As noted previously, the question of how the beat interval is initially inferred such that it can be used to inform SMA falls outside the scope of this paper: see Outstanding Questions.)

The beat-anticipatory trajectory in SMA plays the role of the timing component of the “simulated action” posited by the ASAP hypothesis. Note, however, that it need not correspond to a specific action. We will refer to a beat-anticipatory trajectory in SMA as a “proto-action” because this type of brain activity appears to underlie and temporally structure action without specifying a particular physical movement.

### Anticipating sound and correcting timing

The proposal that neural trajectories in SMA keep time between beats is enriched by incorporating another line of SMA research that attributes it a role in predicting and canceling the sensory consequences of one’s own actions, particularly in the auditory domain [49–51]. This allows an animal to focus on unexpected, i.e., informative, acoustic events. In the language of motor control, SMA is thought to provide an **“efference copy”**of motor commands that can be used by a “forward output model” [52] to predict the sensory feedback that will be produced by those commands. The function of a forward output model has traditionally been ascribed to cerebellum [53–55], which is involved in beat perception [7,56]. However, collected evidence, especially from speech research, suggests that SMA may be driving auditory predictions using more flexible forward models in premotor and parietal cortex, via the dorsal auditory pathway [19,27,57,58].

Given this role of SMA, the cyclic dynamic state of SMA during rhythmic movement can be understood as providing an efference copy of cyclic motor commands to a forward predictive model. Importantly, it has been proposed that forward models can also be used for prediction in the absence of movement-generating motor commands [14,59,60]. In this case, the dynamic state of SMA might better be interpreted as providing a sound-predictive model with an estimate of progress through the time course of a temporally structured stimulus. (The “active inference” framework brings these two ideas together by proposing that movement itself is actuated not through “commands” but through processes that represent the anticipated sensory feedback from movement -- see [61,62].)

Thus, we can flesh out our hypothesis by positing that as SMA tracks progress through each beat interval, it informs a selected auditory forward model that dynamically anticipates auditory input over the course of the interval. The predictions of the forward model could be used to attenuate the perception of uninformative sounds or to increase gain on informative sounds as needed [60,63]. In addition to anticipating a sound that often (but not always) marks the position of the beat, a forward auditory model might also learn to predict intervening sounds between beats, e.g., subdivisions of the beat.

The combination of a proto-action timing process in SMA and a forward model predicting auditory input at specific points in time constitutes a “simulated action for auditory prediction” as proposed by ASAP. We propose that cyclic dynamics in SMA informing premotor forward models are the source of the delta-frequency oscillations originating in the motor system that have been proposed to scaffold temporal prediction and active audition [5,21,60,64,65], though we apply the word “oscillation”cautiously, as we discuss below.

A forward model of expected sound also provides a mechanism for aligning trajectories in SMA to the cycle of sound. During covert beat maintenance, the timing of real sounds and sounds predicted by a forward model could be directly compared, and timing errors could be used to adjust SMA’s representation of progress through the cycle. A similar process could help SMA track the progress of an actual movement based on auditory feedback. Such processes of comparison and adjustment are at the heart of the predictive processing/active inference modeling framework we discuss below.

## Dorsal striatum in beat maintenance

Like SMA, dorsal striatum (especially the putamen) is consistently activated by tasks requiring covert maintenance of a beat [8,35,66]. Dorsal striatum seems especially important to top-down (rather than stimulus-driven) aspects of beat perception: it is more activated by beat maintenance than by beat inference [22], and striatal dysfunction specifically impairs tracking of rhythms requiring internal maintenance of a beat [56,67],

### Initiating and seguencing simulated actions

Motivated by the ASAP hypothesis, we look to the role of dorsal striatum in movement to unravel its role in beat-based timing. In the canonical model of the basal ganglia’s role in motor control [68,69], the dorsal striatum serves as the first link in the motor “direct pathway” (Figure 1B). In the leadup to action initiation, motor areas and other cortical and thalamic areas representing sensory and cognitive context send inputs to dorsal striatum, activating populations representing possible actions. These populations compete with each other through mutual inhibition such that only one self-consistent action plan ultimately survives. It is not yet clear how the right time for action initiation is selected, but it is widely hypothesized that a short pulse of dopamine delivered to striatum from dopaminergic neurons in **substantia nigra pars compacta**provides the impetus to initiate action [70,71], This pulse is thought to cause a burst of activity in dorsal striatum that propagates through the direct pathway (from striatum to globus pallidus to thalamus to cortex, see Figure 1B) leading to the disinhibition of populations corresponding to the selected action in motor and supplementary motor areas [68],

Motor neuroscience draws a distinction between “externally triggered” actions, where action timing is determined by a sensory cue, and “self-initiated”/“internally cued” actions, where action timing is determined by an internal process. Numerous studies indicate that SMA is essential for establishing the timing of self-initiated actions, and that the interaction of SMA and dorsal striatum plays a key role in self-initiated (but not externally-triggered) movements [72–78], It seems that when there is no external cue, the input from SMA to dorsal striatum is essential for activating the direct pathway and initiating action.

Building on these findings, we hypothesize that in covert beat maintenance the direct pathway is using input from SMA to striatum as an internal cue to initialize the next proto-action in SMA, just as it draws on SMA input as a cue to initialize self-generated action during actual movement. This is accomplished by disinhibiting the appropriate sub-population in SMA to generate the neural dynamics that will unfold over the course of the next beat cycle. (See [79], which similarly proposes that the basal ganglia disinhibits selected “emulations” of action for prediction.) The role of striatum in detecting and responding to the state of a cortical timer hearkens back to the influential “striatal beat frequency” model, which proposed that striatum detects a specific state of an ensemble of cortical oscillators as a cue that a specific duration has elapsed [80]. However, our proposal differs in two respects: first, that the cortical input is already tuned to represent a specific time course, and second, that the striatum serves not just to detect a cortical state but to initiate a new process in response.

The importance of striatal dopamine in initiating self-generated actions suggests that dopamine may also play an important role in sustaining an internal representation of a beat. See Box 1 for discussion. Further, the position of the striatum at an interface of anticipatory timing and movement suggests possible neural substrates for the link between beat-based timing and movement, discussed in Box 2.

#### BOX 1: Dopamine and beat percept strength

The neurotransmitter dopamine is widely associated with reward anticipation and motivation. However, these associations are most appropriate to dopamine in the cortex and ventral striatum, provided by dopaminergic neurons in the midbrain’s ventral tegmental area. By contrast, dopamine in dorsal striatum, delivered by dopaminergic cells in the nearby substantia nigra pars compacta (“nigrostriatal dopamine”), is more closely associated with (and capable of triggering) the initiation of action [118]. It is possible that nigrostriatal dopamine is similarly involved in initiating the time-keeping proto-actions in SMA that we have discussed above.

A current line of research suggests that dopamine (and nigrostriatal dopamine in particular) may encode certainty in cognitive constructs. Friston et al. [119] argue that “dopamine reports the precision or salience of sensorimotor constructs (representations) encoded by the activity of the synapses they modulate,” and therefore that “changing the levels of dopamine changes the level of uncertainty about different representations”[119]. One such “sensorimotor construct” could be an internal representation of the period and phase of a temporal cycle underlying an ongoing sound. In this case, dopamine would also serve as an index of confidence in temporal predictions based on that cycle. Recent experimental evidence has offered support for this interpretation [120–122] in particular, striatal dopamine level has been found to index temporal expectation of perceptual cues [123] and certainty about timing [124],

Combining these ideas with our current framework, we hypothesize that when SMA accurately anticipates rhythmic timing, striatal dopamine levels rise, making it easier for the striatum to initiate the proto-actions that serve as an internal representation of the beat cycle and strengthening the internally generated component of the beat percept.

#### BOX 2: Insight into dance and Parkinson’s

The role of the basal ganglia in selecting, sequencing, and disinhibiting movement, combined with their proposed role in doing the same for time-keeping proto-actions suggests that this may be the key point of intersection for musical anticipation and movement. Several processes in the basal ganglia have global motor-modulatory effects that are not limited to specific movements, and that therefore may be responsible for our inclination to move (and our facility at moving any body part) along with rhythmic music. The pulses of striatal dopamine from the substantia nigra that are thought to time the output from dorsal striatum could have the effect of encouraging initiation of motor activity at the same moments as the next proto-action is initiated in SMA. Action initiation also appears to require overcoming the anti-kinetic effect of the “hyperdirect” pathway through basal ganglia [125,126]. If the hyperdirect pathway’s influence is attenuated to allow for striatal output initiating the next forward model, this may have the additional effect of broadly facilitating the initiation of movement.

Both of these mechanisms -- dopaminergic and hyperdirect -- are plausible links to the remarkable gaitfacilitating effects of rhythmic cueing in Parkinson’s [127], An extra dopamine pulse could help overcome the dopamine deficiency thought to make action initiation so difficult in Parkinsons; alternatively (or additionally), a drop in hyperdirect pathway influence could make it easier for weak dopaminergic signals to successfully initiate action. And both mechanisms offer a possible mechanism for the modulation of beta (~2OHz) oscillation power that has been repeatedly observed to synchronize with a beat [128,129]: the anti-kinetic effect of the hyperdirect pathway is linked to beta [130,131], and phasic striatal dopamine is thought to attenuate beta [132],

### From action sequencing to metrical anticipation

To anticipate auditory input in a uniformly repeating cycle (e.g., Figure 1A), the basal ganglia could select the same pattern of activity in SMA over and over again. Yet why should such a complex brain structure be necessary to play this role -- surely the flexible dynamics of SMA could sustain a repeating cycle on its own? Crucially, cyclic movement and musical rhythm both have repeating temporal structure at multiple time scales. In cyclic movement, multiple actions are sequenced into longer cycles such as the left-right cycle of stepping, and in music, metrical structure contains repeating temporal patterns at multiple scales such as beats, groups of beats, etc. For a brain structure like SMA that seems to specialize in generating dynamics over subsecond time scales, flexibly structuring “chunks” of dynamic activity into longer repeating patterns may not be trivial.

Thus, the motor function of dorsal striatum most relevant to our sense of the beat may be its role in sequencing movements. Striatal activity is important for making transitions between sequential motor “chunks” in learned motor sequences, both in mice [81–83] and humans [84], During rhythmic action, striatal activity (in monkeys) differs predictably from one action repetition to the next [85], suggesting that repetitions of superficially similar movements are being represented as a sequence of distinct actions in striatum. Sequential activity in striatum during covert timing tasks (in mice) suggests that even undifferentiated wait time is represented as a sequence of distinct states [86]. Models suggest that sequential transitions in striatum may be orchestrated by patterns of inhibitory connectivity among striatal neurons trained by sequential motor cortical input [87], or else by a “competitive queuing” process among cortical representations [88,89] (a mechanism recently used to model rhythmic sequencing [90]).

Following this thread of evidence, we hypothesize that striatum sequences the short proto-actions of SMA into longer repeating patterns. Each beat in, for example, group of 4 (a common repeating unit in Western music) would have its own characteristic identity in SMA as a proto-action with identical duration but different signatures in neural dynamics. As one reached its end, the dorsal striatum would initiate the next in the sequence (see Figure 3). Thus, activity in both SMA and striatum should show patterns that repeat with the metrical cycle as well as with the beat; this is consistent with a recent EEG-fMRI study [91]. Recent observations in rodents suggest that striatum orchestrates hierarchically-as well as sequentially-structured movement [92] and thus might support hierarchical metrical structures [93].

**Figure 3:**
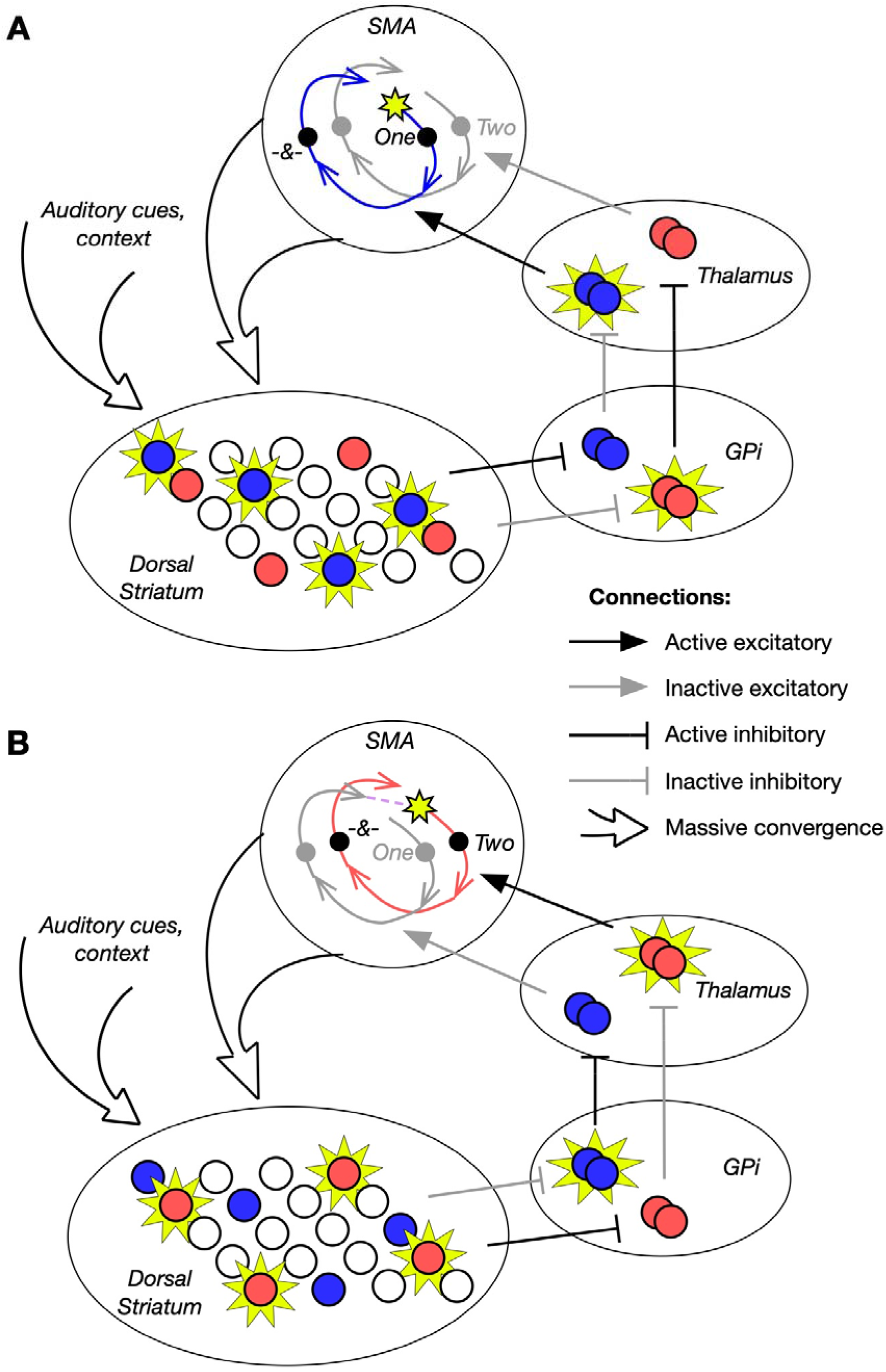
Basal ganglia sequences the beat-tracking trajectories in SMA. A) Once a beat has been inferred in an auditory rhythm, neural firing rates in SMA follow a trajectory which tracks progress through a beat cycle, informing the anticipation of the beat and possibly other sounds locked to the cycle (such as a subdivision of the beat, indicated by the “&” symbol on the trajectory). The activity of neurons generating this trajectory is initiated and reinforced by selective disinhibition by the basal ganglia: a population specific to the first beat cycle is active in dorsal striatum (blue). Acting through the internal segment of the Globus Pallidus (Gpi), this population disinhibits a thalamic population, which provides SMA with excitation specific to that trajectory. B) As the first beat cycle ends and the second beat arrives, some combination of SMA’s evolving input to dorsal striatum and inhibitory interactions within dorsal striatum activates a new striatal subpopulation (red). This population disinhibits a new subpopulation in SMA that allows SMA activity to continue to evolve along a new trajectory similar to the previous one. It spans the same time interval, but plays a different role in the metrical structure and may inform different auditory expectations specific to that beat cycle.

An advantage of a framework that separates timing and sequencing activity into separate brain regions is that it allows for metrical structures that include non-isochronous durations as “beats” [94] as in various traditional musics like Balkan and Scandinavian music, where such musical structures inspire tightly coordinated rhythmic movement in enculturated performers and dancers [95–97], For example, each grouping in a “2/2/3” Balkan pattern, in which every third inter-beat-interval is 1.5 the duration of the previous two intervals, would be represented by its own distinct proto-action in SMA and sequenced by dorsal striatum, with the third proto-action being longer in duration than the other two.

## Implications for beat perception modeling

The covert beat-maintenance process we hypothesize based on the neurophysiology of motor processing can be loosely summed up as follows. An adjustable-speed neural process (“proto-action”) in SMA aligns to perceptual intervals, informing auditory anticipation over the course of the interval and correcting its alignment based on continuing auditory cues. On each beat cycle, output from SMA signals dorsal striatum to select and disinhibit a new cycle-specific neural subpopulation in SMA to generate the next proto-action. In combination, when the proto-actions sequenced by striatum and executed by SMA are of a consistent duration, this creates a self-sustaining cycle of internal activity that creates a beat-based pattern of auditory anticipation.

This system can be understood as a neurophysiological implementation of the “attending rhythms” of dynamic attending theory [98,99] that are posited to direct attention overtime. However, our neurophysiological hypotheses are only partially consistent with the oscillator-based models applied by dynamic attending theory to beat perception [100–102], Since our proposed brain dynamics do not account for beat inference, they may be consistent with a model in which beat inference relies on a bank of oscillators [101]. However, describing cyclical activity in SMA as a limit cycle oscillation during beat maintenance suggests that the process is an autonomous, homogeneous cycle like a pendulum, whereas we hypothesize that active input from the striatum shapes this activity cycle by cycle. Notably, our account allows for a simpler model of the generation of non-isochronous beat patterns than is possible with homogenous oscillators. We also believe our account generalizes more easily than an oscillator account to non-isochronous rhythmic anticipation in speech [103–105]. The sequential nature of our neurophysiological account of the beat would be better described by the model of concatenated diffusion processes proposed and evidenced by Merchant& Averbeck [106].

The idea of SMA dynamics aligning to the beat based on a forward model of expected sound is closely related to the “predictive processing”/“active inference”school of brain modeling. According to this perspective (recently invoked in describing and modeling musical rhythm [107–109]), the brain is constantly predicting its inputs and adjusting its representations of states in the world based on its errors in prediction. In our picture of SMA, a dynamic neural state represents either physical progress through a cyclic action or progress through an externally-generated cycle of auditory events. In either case, predictions of auditory input based on this dynamic state are compared to actual auditory input, and errors are used to correct the state. We are developing a computational model that expresses this process in terms of Bayesian inference, compatible with predictive processing [110]. This will help connect our neurophysiological picture to Bayesian models of timing (reviewed in [111]; see also [112], which connects Bayesian timing inference with SMA dynamics).

## Concluding remarks

Beat perception offers cognitive science an exciting opportunity to explore how cognition and action are intertwined in the brain even in the absence of movement. Based on the idea that simulated actions support the precise temporal predictions involved in covert beat maintenance, we have drawn on the motor neuroscience literature to propose specific roles for SMA and the dorsal striatum, two key motor structures engaged by beat perception. This leads to testable predictions, listed below, and illustrates how cognitive science, by engaging with the details of motor neuroscience, can move toward biologically-detailed models of embodied cognition.

Our hypotheses lead to several testable predictions in humans. First, if dorsal striatum is engaged in sequencing beats into higher-order metrical structures, then these slower metrical periodicities should be reflected in striatal activation, which, unlike the beat cycle itself, may be slow enough to directly observe in fMRI [113]. Second, the hypothesis that each beat cycle represents a distinct simulated action suggests that characteristic beat-related activity in scalp EEG recordings should align to and anticipate non-isochronous beats in enculturated listeners, e.g., in Balkan music [95,96]. Third, recordings in human basal ganglia (e.g., during deep brain stimulator implantation) should show activity at a characteristic phase of the beat cycle, regardless of the specific auditory events implying the presence of a beat. Fourth, this activity should include striatal dopamine pulses that may be observable using fastscan cyclic voltammetry [114], which should also detect elevated tonic striatal dopamine in conditions of a strong internal beat (see Box 1). Finally, ECoG recordings in human SMA should show firing rate trajectories aligning to the beat cycle during complex rhythms and accounting for variation in performance on beat-based timing tasks, and perturbations of these trajectories [115] should disrupt beat maintenance.

## Acknowledgements

We are grateful to Nancy Kopell, Michelle McCarthy, Jeffrey Markowitz, Sean Patrick, and our reviewers for their input and insights.

## Outstanding Questions

⍰ **How do the proposed mechanisms of beat maintenance interact with the larger network of brain regions engaged by beat-based stimuli?** The cerebellum is known to be active in beat-based timing tasks. It plays a role in rapidly relaying temporally precise information to frontal areas, and thus could be an important input to the beat maintenance circuit. It also contributes to error-correction in motor control, and could similarly serve to update beat phase and tempo in response to errors in anticipation. Further, we have only touched briefly on the bidirectional interaction between the beat maintenance system, premotor areas, and the auditory cortex. This interaction is central to beat perception and should be more fully characterized.
⍰ **How does the brain perform beat inference?** Before the beat maintenance system can be engaged, the presence of a beat must first be recognized, a process that is likely inseparable from initial inference of its tempo and phase. Beat inference has been modeled as the emergence of resonance in a bank of oscillators. It could also involve a process of characterizing temporal structures and detecting their repetition, possibly utilizing coincidence detection between internal timing signals and auditory events in cerebellum and/or basal ganglia. To transition from beat inference to beat maintenance, a tempo must be persistently represented in the brain, as evidenced by our capacity to continue a beat after the end of a rhythm. One logical place to seek this representation would be the dorsolateral prefrontal cortex, a region that supports persistent representations and is specifically activated by tempo-based tasks.

- **How does the development of beat perception relate to motor system development?** Neural research suggests beat perception is present in newborns, prior to the emergence of stable motor synchronization to a beat (around age 3-5). Further work is needed to determine whether infants accurately predict beat timing using their motor system.

## Glossary

Action Simulation for Auditory Prediction (ASAP) hypothesis: Posits that the beat underlying a rhythmic auditory stimulus is tracked and anticipated by repeated movement-like processes in the motor system that are coordinated through rapid bidirectional communication with auditory cortex.
Beat inference: Recognizing that a rhythm supports a beat and inferring its tempo and phase.
Beat maintenance: Implicit anticipation of the timing of each beat underlying a rhythmic auditory stimulus.
Dynamical systems: Refers to systems of variables whose direction of change is determined by their present state, and to the field of mathematics that studies such systems.
Efference copy: A copy of a neural signal that directs movement, sent to sensory processing areas to anticipate the sensory consequences of the movement.
Functional magnetic resonance imaging (fMRI): A technique for measuring brain activity by detecting the changes in blood oxygenation and flow that occur in response to neural activity.
Isochronous: Occurring periodically in time with a constant interval between events.
Proto-action: A term coined here to describe a neural process that provides the temporal structure of a movement but is not specific to one movement type or body part.
Putamen: The subregion of striatum most associated with motor function.
Striatum: The main input structure to the basal ganglia.
Substantia nigra pars compacta: A structure in the basal ganglia formed by dopaminergic neurons. The loss of these neurons is characteristic of Parkinson’s disease.
Supplementary motor area (SMA): The caudal subregion of the medial premotor cortex. Also called “SMA proper” in some sources to disambiguate from neighboring “pre-SMA.” Contributes to motor control, planning, and coordination.
Tonic input: An input to a neural process that does not change over time.

